# Single cell profiling reveals GSM-15606 attenuates air pollution-induced inflammation and preserves hippocampal neurogenesis

**DOI:** 10.64898/2026.01.27.702094

**Authors:** Mohammad Shariq, Wanqing Pan, Xiran Chen, Wang Xiang, Jose Godoy-Lugo, Lei Peng, Jonathan N. Levi, Albina Ibreyeva, Kristina Shkirkova, Wynnie Nguyen, Constantinos Sioutas, William J. Mack, Caleb E. Finch, Max A. Thorwald, Michael A. Bonaguidi

## Abstract

1.

**Introduction:** Air pollution (AirPoll) is a major environmental risk factor for age-related cognitive decline and dementia, yet we poorly understood the cellular and molecular mechanisms underlying its effects and their potential attenuation.

**Methods:** We combined single cell RNA sequencing with immunohistochemistry to determine transcriptional responses in microglia, astrocytes, neurons and neural stem cells in the hippocampus of mice following exposure to chronic diesel exhaust particle (DEP). Differential gene expression profiles were compared between filtered-air and DEP exposed animals. The gamma secretase modulator GSM-15606 (BPN) was used to probe selective rescue of inflammatory signatures across distinct cell populations.

**Results:** DEP exposure triggered robust inflammatory programs in microglia and astrocytes, including upregulation of cytokine signaling components, innate immune receptors, stress-responsive transcription factors, and markers of reactive glial phenotypes. In neural stem cells, DEP induced activation of gliosis-associated pathways, including Il6st, Stat3, and Txnip, consistent with a pro-inflammatory state that may bias lineage decisions. Immunostaining confirmed a significant reduction in immature neurons in the neurogenic niche after AirPoll exposure. GSM-15606 attenuated many DEP-induced transcriptional alterations in microglia and astrocytes, reducing expression of inflammatory mediators and reactive gliosis markers, but did not modulate the inflammatory profile of neural stem cells.

**Conclusions:** AirPoll activates divergent inflammatory pathways across hippocampal cell populations and suppresses neurogenesis. Targeting inflammation with GSM-15606 selectively reverses glial but not neural stem cell responses, highlighting cell-type-specific mechanisms and potential therapeutic pathways to mitigate pollution-related cognitive vulnerability. These results support GSM-15606 as a protective agent against AirPoll-induced hippocampal dysfunction and amyloidogenic stress.

## 2. Background

AirPoll is recognized as a major contributor to age-related cognitive decline and dementia. Several large observational studies show that long-term exposure to fine particulate matter (PM_2.5_) and traffic-related pollutants is associated with higher rates of Alzheimer’s disease (AD) and earlier cognitive deterioration (Wilker et al., 2023; Shi et al., 2020; Mork et al., 2023). Notably, population-level improvements in air quality, such as those following China’s Clean Air Act, correspond with measurable benefits in cognitive performance among older adults (Yao et al., 2022). These findings align with mechanistic work in humans and animal models showing that pollutants trigger chronic neuroinflammation, oxidative injury, blood-brain barrier dysfunction, and widespread activation of microglia and astrocytes, all of which overlap with early features of AD pathology (Calderón-Garcidueñas et al., 2008; Cory-Slechta & Sobolewski, 2023; Iaccarino et al., 2021). A growing number of studies also indicate that particulate exposure disrupts adult hippocampal neurogenesis, a form of brain plasticity important for learning and memory (Boda et al., 2020; Jaiswal & Singh, 2024; Woodward et al., 2018). Both traffic-related air pollutants (TRAP) and PM exposures reduce proliferating progenitors, decrease NeuN/BrDU-positive newborn neurons, and promote reactive gliosis within neurogenic regions(Costa et al., 2017; Jaiswal & Singh, 2024; Kilian et al., 2023; Sahu et al., 2021; Woodward et al., 2018). Diesel exhaust particles (DEP) and nanosized PM (nPM) induce similar effects, together with pronounced microglial activation and synaptic vulnerability in hippocampal circuits(Cheng et al., 2017; Levesque et al., 2011; Morgan et al., 2011; Zhang et al., 2023); Godoy-Lugo et al, 2025). Although PM_2.5_ has historically been the most common exposure model, variability in its composition (Thorwald & Finch) has led many groups to use standardized DEP preparations, which offer more reproducible and robust neuroinflammatory outcomes (Block & Kodavanti, 2021; (Shkirkova et al., 2024) Farahani et al, 2021).

We recently reported (Godoy-Lugo et al., 2024) that the γ-secretase modulator GSM-15606 (BPN) decreased brain levels of the Aβ42 peptide in mice exposed to 8 weeks of nano-scale nPM or DEP. GSM-15606 is known to reduce glial reactivity and improve several AD pathological features without disrupting Notch and other essential signaling pathways (Wagner et al., 2017; Rynearson et al., 2021; Prikhodko et al., 2020). We do not know whether GSM-15606 benefits extend to the AirPoll inflammation, especially within the vulnerable neurogenic populations of the hippocampus. This study used single cell transcriptomic analysis to examine how GSM-15606 influences the cellular and molecular responses of these neurogenic populations following AirPoll exposure. This approach allows us to determine whether a brain-penetrant compound can reduce the inflammatory and gliotic changes that impair neurogenesis and may offer a clearer understanding of how therapeutic interventions could ameliorate air pollution-induced brain damage.

## 3 Methods

### 3.1 Animals

All animal procedures were approved by the Institutional Animal Care and Use Committee at the University of Southern California and conducted under the oversight of the USC Department of Animal Resources (#20842). Mice on a C57BL/6 background were housed in groups of four under controlled environmental conditions at 22°C with 30% relative humidity and maintained on a 12-hour light–dark cycle with standard bedding and nesting materials. The animals used in this study were wild-type and carried a Nestin transgene (Jackson Laboratory, stock no. 034387), which was utilized for cell isolation in complementary hippocampal analyses.

### 3.2 Treatment and diesel exhaust particle exposure

Male and female mice were administered the γ-secretase modulator GSM-15606 ((S)-N-(1-[4-fluorophenyl]ethyl)-6-(6-methoxy-5-[4-methyl-1H-imidazol-1-yl]pyridin-2-yl)-4-methylpyridazin-3-amine benzenesulfonate, besylate salt) at a dose of 10 mg/kg/day, with vehicle control. GSM-15606 was incorporated into standard chow libitum beginning one week prior to exposure and continued throughout the eight-week exposure (see Figure S1). Mice were subjected exclusively to diesel exhaust particles (DEP), a well-characterized component of TRAP PM_2.5_, or filtered air control in two independent cohorts that were studied at separate time points. DEP exposure was conducted at a concentration of 100 μg/m^3^ for 5 hours per day over a period of 8 weeks. Standardized DEP material was obtained from the National Institute of Standards and Technology (NIST; SRM 2975). For aerosol generation, DEP was suspended in Milli-Q deionized water, thoroughly mixed, and sonicated for 30 minutes to ensure uniform particle dispersion prior to re-aerosolization for animal exposure.

### 3.3 Single cell RNA-seq processing and quality control

Raw base call files from 10x Genomics libraries were processed with Cell Ranger (v8.0.1) using the default gene expression pipeline and alignment to the mm10 reference genome to generate cell-by-gene UMI count matrices. Subsequent analyses were performed in R (v4.4.3) using Seurat (v5.3.0) and custom scripts. Low-quality barcodes were excluded based on library complexity and mitochondrial content: cells with >10,000 detected genes (nFeature_RNA) or >25% mitochondrial RNA (percent.mt) were removed, and the remaining 6,916 cells (23,812 detected genes across the dataset) were retained for downstream analyses. Counts were normalized using Seurat’s NormalizeData function (LogNormalize; scale factor = 10,000) and stored in the assay data slot. Highly variable genes were identified using FindVariableFeatures (top 3,000 features), followed by scaling with ScaleData. Principal component analysis (PCA) was performed on the highly variable gene set, and the number of components retained for downstream steps was guided by inspection of elbow plots and JackStraw analysis.

### 3.4 Dimensionality reduction, clustering, and cell-type annotation

A shared nearest neighbor (SNN) graph was constructed using FindNeighbors on the top principal components (dims = 1:20), and cells were clustered using the Leiden algorithm (FindClusters, resolution = 0.2). Nonlinear dimensionality reduction was performed with UMAP (RunUMAP) for visualization of global transcriptomic structure. Cluster identities were assigned based on expression of canonical marker genes using FindAllMarkers, inspection of UMAP feature plots (FeaturePlot), and comparison with published hippocampal and neurogenic niche datasets (Hart J, Harris L (2025), Chenxu Chang, Hongyan Zuo, Yang Li (2023). Major classes included neural stem and intermediate progenitor cells (NSC/IPC), neurons, oligodendrocytes, microglia, astrocyte-lineage populations, pericytes, and endothelial cells.

### 3.5 Differential expression analysis

Differential expression analyses were performed using edgeR (v4.6.3). For each comparison, cells from the cluster of interest were subset by condition, and raw UMI counts were extracted from the Seurat object (GetAssayData, slot = “counts”). Genes were retained if they showed nonzero mean expression in at least one group, and the filtered count matrix was used to construct an edgeR DGEList. Library sizes were normalized using the trimmed mean of M-values (TMM) method (calcNormFactors), dispersions were estimated with estimateDisp, and a quasi-likelihood negative binomial generalized linear model was fit with glmQLFit using a design matrix of ∼group. Differentially expressed genes were identified using quasi-likelihood F-tests (glmQLFTest), and p values were adjusted for multiple testing using the Benjamini-Hochberg procedure (FDR).

### 3.6 Gene ontology enrichment

Gene Ontology (GO) enrichment analysis was performed using clusterProfiler (v4.16.0) with mouse gene annotations (org.Mm.eg.db). Differentially expressed genes were separated by direction of change (upregulated and downregulated) prior to enrichment analysis. For each gene set, over-representation testing was conducted with enrichGO using gene symbols as identifiers (keyType = “SYMBOL”), Benjamini-Hochberg adjustment (pAdjustMethod = “BH”), and a p-value cutoff of 0.05; qvalueCutoff was set to 0.99. GO results were computed across ontologies (ont = “ALL”) and summarized using standard clusterProfiler visualization workflows.

### 3.7 Gene set scores

Gene set scores were computed for inflammatory de-activation with GSM-15606 in the AirPoll context. This gene set is composed of genes that are inflammation-related and are upregulated in AirPoll. Module scores were visualized using Seurat VlnPlot and overlaid boxplot (center line = median; box = interquartile range; whiskers = 1.5xIQR; outliers hidden). Pairwise differences in module scores across conditions were assessed using two-sided Wilcoxon rank-sum tests implemented via ggpubr::stat_compare_means with predefined comparisons (Air_pollution_plusGSM vs Air_pollution, Air_pollution vs Filtered_air, Air_pollution_plusGSM vs Filtered_air, Filtered_air_plusGSM vs Filtered_air).

## 4 Results

### 4.1 Single-cell characterization of the hippocampal neurogenic niche across exposure and treatment conditions

To define how (DEP exposure alters the hippocampal neurogenic niche and whether the gamma-secretase modulator GSM-15606 mitigates these effects, we assigned 16 Nestin::CFP mice to four treatment groups (n = 4 per condition: filtered air, filtered air + GSM-15606, DEP exposure, and DEP exposure + GSM-15606). Treatments began at 4 months of age, and mice were sacrificed at age 6 months for 10x Genomics single-cell RNA-seq and immunohistochemistry (Figure 1A). Raw 10x Genomics single-cell RNA-seq libraries were processed with Cell Ranger, followed by a single, globally applied QC workflow across the merged dataset. Cells were retained if mitochondrial transcript fraction was ≤25% and if the number of detected genes was ≤10,000, yielding a final dataset of 4,853 cells spanning 23,812 detected genes (Supplementary Figure 1).

**Figure 1.**
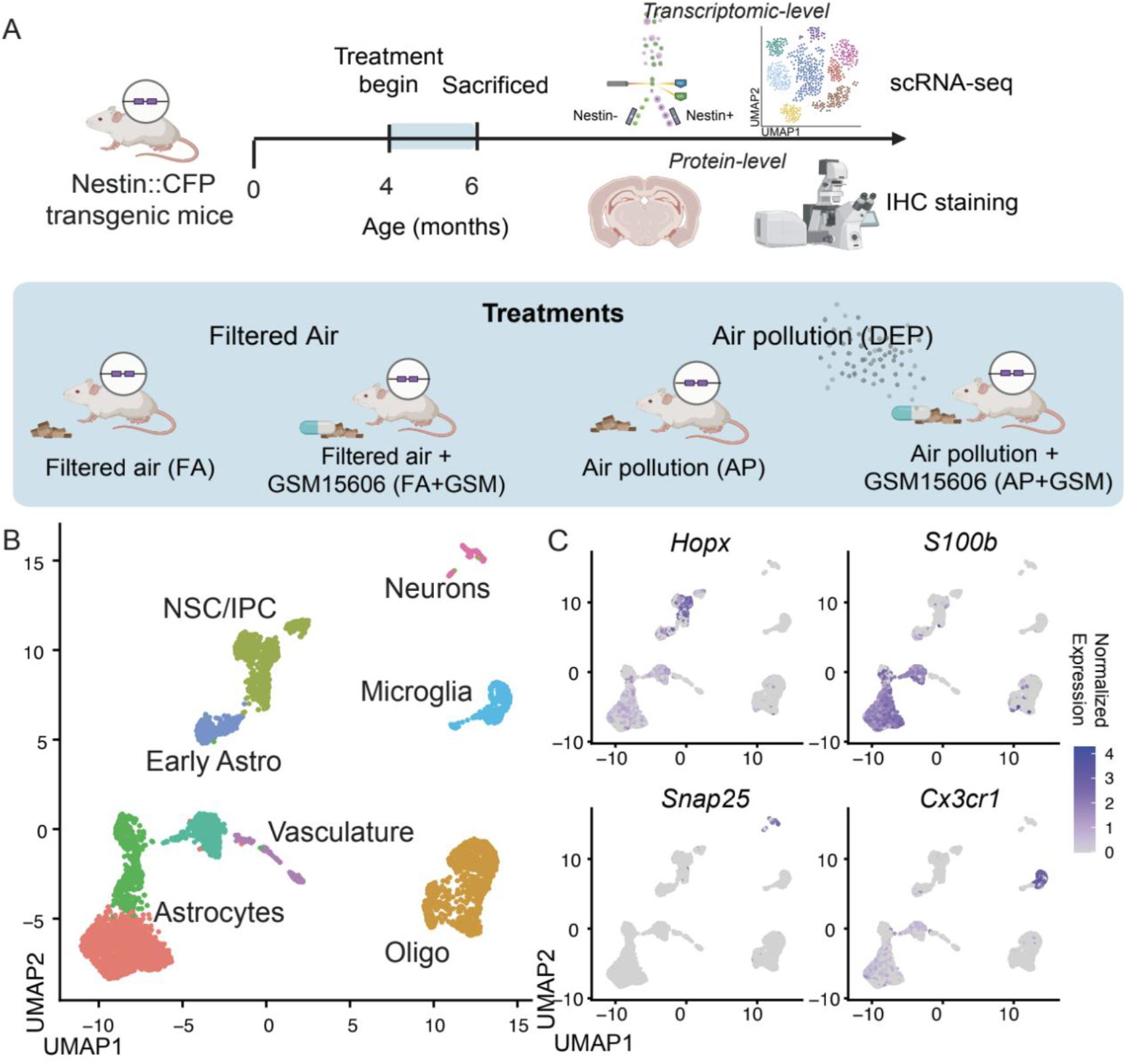
Transcriptomic overview of the Nestin-lineage hippocampal cell type response to air pollution and GSM-15606 treatment. (A). Experimental schematic showing Nestin::CFP transgenic mice exposed to filtered air, diesel exhaust particle (DEP) air pollution, or DEP with GSM-15606 treatment, followed by tissue collection for FACS sorting of Nestin-positive cells for scRNA-seq and immunohistochemistry (IHC). (B). UMAP embedding of all captured cells colored by major annotated populations, including Neural Stem Cells and Intermediate Progenitor Cells (NSC/IPC), microglia, astrocyte-lineage clusters (including early Astros), oligodendrocytes (Oligo), neurons, endothelial cells and pericytes (vasculature). (C). Feature plots of canonical markers used for cluster annotation, highlighting NSC/IPC (*Hopx*), astrocyte-lineage (*S100b*), neurons (*Snap25*), abd microglia (*Cx3cr1*).

Unsupervised analysis in Seurat identified the major cellular classes of the hippocampal neurogenic niche. Briefly, after normalization, variable-feature selection, scaling, and PCA, cells were embedded with UMAP and clustered using Leiden community detection, resolving neural stem/progenitor populations, neurons, oligodendrocytes, microglia, astrocyte-lineage populations (including an early astrocyte-like population), and vascular-associated populations including endothelial cells and pericytes (Figure 1B--C; Supplementary Figure 2). Cell-type composition summaries are shown in Supplementary Figure 3 and were used to contextualize downstream analyses aimed at distinguishing transcriptional programs from shifts in cellular abundance.

### 4.2. Microglia and astrocytes display unique inflammatory signatures in response to DEP exposure

Single-cell RNA profiling revealed a pronounced microglial response to diesel exhaust particle exposure. Microglia formed a well-defined cluster, enabling targeted differential expression analysis between filtered air and pollution conditions (Fig 2A). Following exposure, microglia showed significant upregulation of canonical inflammatory mediators, including Il1b and Cxcl2, which are central drivers of cytokine and chemokine signaling in activated microglia and have been consistently linked to particulate-induced neuroinflammation (Levesque et al., 2011; Costa et al., 2017). Expression of Lgals3, a marker associated with microglial reactivity and phagocytic remodeling, was also increased, consistent with reports of pollution-driven microglial activation states (Zhang et al., 2023).

**Figure 2.**
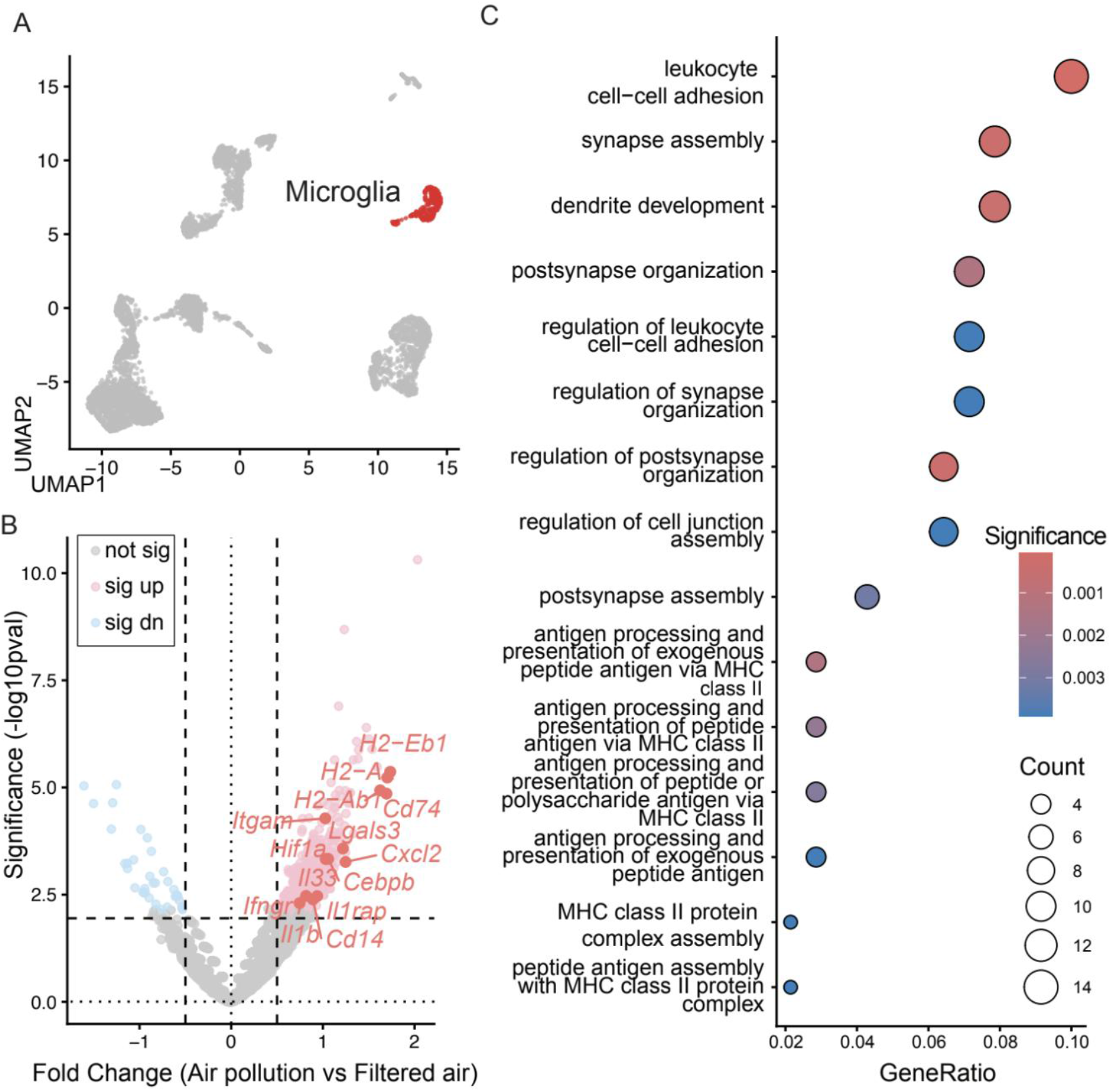
Microglia activate an air-pollution–induced inflammatory program. (a) UMAP highlighting the microglia cluster within the full dataset. (b) Volcano plot of microglial differential expression for air pollution (DEP) versus Filtered Air, with AP-upregulated inflammatory genes annotated. (C) Gene Ontology (GO) enrichment analysis of the top 100 AP-upregulated microglial genes showing over-represented immune/inflammatory biological processes (dot size = gene count; color = adjusted p-value).

In parallel, transcriptional regulators that amplify inflammatory programs under hypoxic and stress conditions, including Hif1a and Cebpb, were significantly elevated. Induction of these factors has been reported in both human and experimental models of AirPoll exposure and is known to reinforce sustained microglial activation (Calderón-Garcidueñas et al., 2008). AirPoll further enhanced expression of innate immune and inflammatory signaling components such as Itgam (CD11b), Cd14, Il1rap, Il33, and Ifngr1, reflecting heightened responsiveness to inflammatory cues and interferon signaling, pathways previously implicated in pollution-associated microglial priming (Costa et al., 2017; Cory-Slechta & Sobolewski, 2023) (Fig 2B).

Notably, several genes encoding major histocompatibility complex class II molecules (H2-Ab1, H2-Aa, H2-Eb1, Cd74) were among the most strongly upregulated transcripts (Fig 2B). Induction of this antigen-presentation machinery is a hallmark of chronically activated microglia and has been observed in models of particulate exposure and Alzheimer’s disease–related neuroinflammation (Sahu et al., 2021; Iaccarino et al., 2021). Gene ontology enrichment analysis of differentially expressed microglial genes further supported this shift, revealing strong overrepresentation of pathways related to antigen processing and presentation, peptide loading, and immune cell interaction (Fig 2C).

In addition to immune-related pathways, enriched gene sets associated with synapse organization, postsynaptic structure, and dendritic remodeling were observed (Fig 2C). These findings align with prior evidence that pollution-activated microglia influence synaptic integrity and neuronal connectivity, linking microglial inflammation to functional vulnerability within hippocampal circuits (Morgan et al., 2011; Kilian et al., 2023). Together, these data demonstrate that diesel exhaust particle exposure drives a robust inflammatory and antigen-presenting transcriptional program in microglia, accompanied by enrichment of pathways associated with immune activation and synaptic remodeling. This result positions microglia as a primary cellular responder to AirPoll within the hippocampus and a likely contributor to pollution-associated cognitive vulnerability.

### 4.3. Air pollution reprograms neural stem cells toward stress-responsive and gliotic states

Analysis of the neural stem and intermediate progenitor cell compartment revealed that diesel exhaust particle exposure alters intrinsic transcriptional programs linked to inflammatory signaling and lineage bias. Differential expression analysis showed increased activity of pathways associated with cytokine responsiveness and cellular stress rather than overt immune activation (Fig 3C). Notably, expression of Il6st (gp130) and Stat3 was elevated in NSCs following pollution exposure, indicating engagement of IL-6 family signaling within the stem cell population (Fig 3B). Activation of this signaling axis is known to influence fate decisions during injury and inflammation by favoring astroglial differentiation and reinforcing gliotic programs (Nakashima et al., 1999; Levesque et al., 2011; Costa et al., 2017). (Fig 3B)

**Figure 3.**
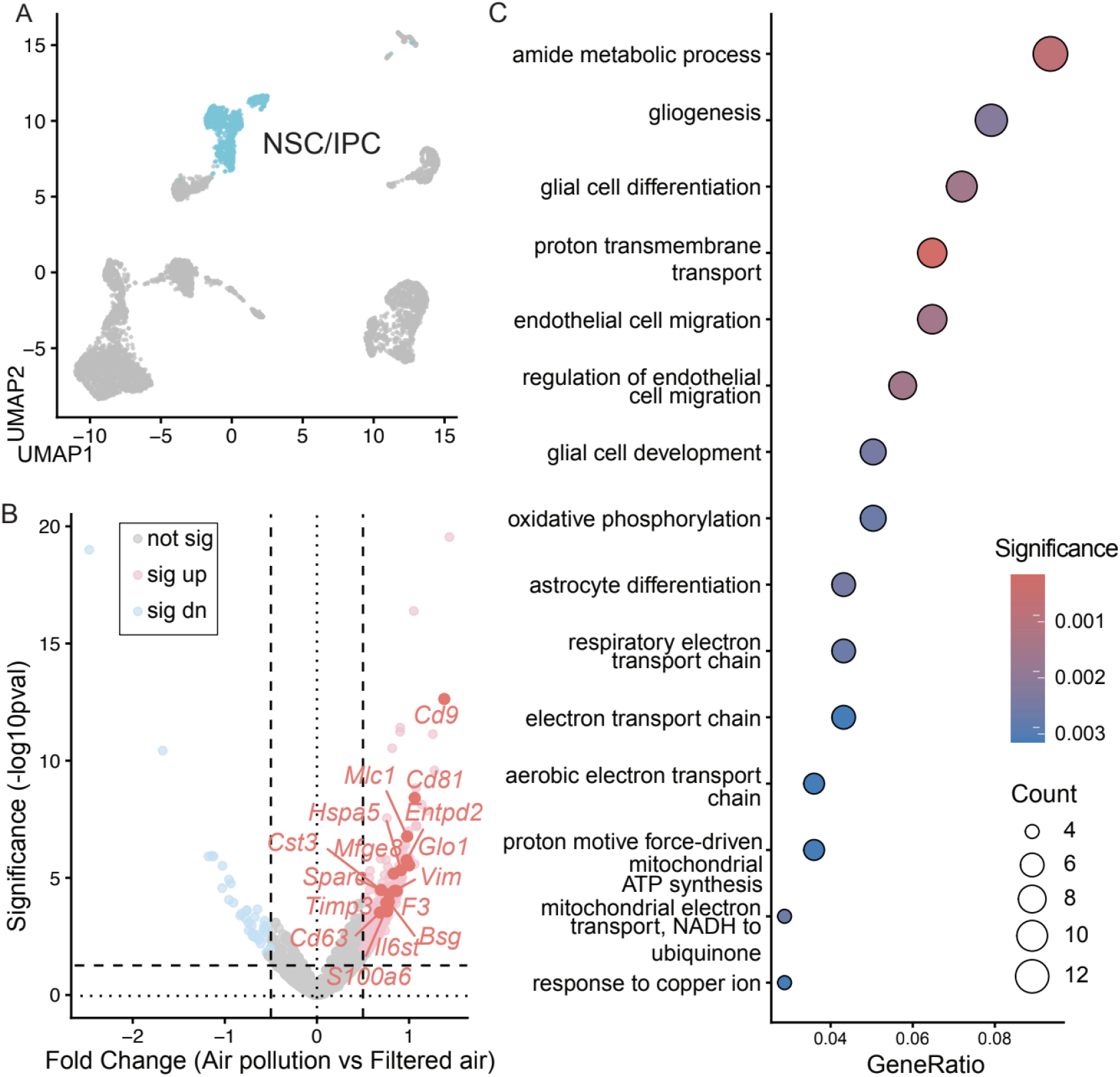
NSC/IPC show a gliosis-like stress response to air pollution. (A) UMAP highlighting the NSC/IPC cluster within the full dataset. (B) Volcano plot of NSC/IPC differential expression for air pollution versus filtered Air, with representative air pollution-upregulated inflammatory genes annotated. (C) GO enrichment analysis of the top 100 AP-upregulated NSC/IPC genes, indicating activation of stress-associated biological processes.

In parallel, NSCs exhibited increased expression of genes associated with oxidative and metabolic stress. Upregulation of Txnip, a regulator of redox balance and inflammatory signaling, supports the presence of a stress-responsive state that has previously been linked to impaired neurogenesis and glial activation in degenerative and inflammatory contexts (Hartz et al., 2008; Kilian et al., 2023). Rather than reflecting a generalized inflammatory phenotype, this transcriptional pattern suggests selective engagement of signaling pathways that can indirectly shape the cellular composition of the neurogenic niche.

Although antioxidant and cytoprotective responses were also evident, including increased expression of *Nfe2l2* and enrichment of mitochondrial and oxidative phosphorylation pathways (Fig. 3C), these changes likely reflect an adaptive response to pollution-induced cellular stress rather than effective restoration of homeostasis. Similar compensatory antioxidant programs have been reported in other models of particulate exposure, particularly in response to metal-rich air pollutants, but are often insufficient to counter sustained upstream inflammatory and oxidative signaling (Scieszka et al., 2022; Zhang et al., 2021,Shkirkova et al., 2026). Recent work further suggests that metal-containing particulate matter can drive lipid peroxidation and ferroptosis-associated stress programs despite activation of antioxidant pathways, creating a state in which redox defense mechanisms are engaged but ultimately overwhelmed. In this context, the coexistence of stress-response and gliosis-promoting pathways within NSCs points to a transcriptional imbalance rather than recovery.

Consistent with this interpretation, gene ontology analysis revealed enrichment of biological processes related to gliogenesis, astrocyte differentiation, and metabolic remodeling. Together, these findings indicate that AirPoll shifts neural stem cells toward a stress-responsive state that favors astroglial outcomes. Rather than acting solely as passive targets of inflammation, NSCs may actively participate in reinforcing a gliosis-prone environment within the polluted hippocampus, thereby contributing to reduced neurogenic output and increased vulnerability of hippocampal circuits over time.

### 4.4. DEP driven inflammation leads to a loss of immature neurons and an increase in astrocytes in the dentate gyrus

Next, we performed immunostaining to examine how diesel exhaust particle exposure affects cellular outcomes within the hippocampal neurogenic niche. Quantification of DCX-positive immature neurons revealed a clear reduction in their number in DEP-exposed animals compared with filtered air controls (Fig 4A-B). This reduction is consistent with our single-cell transcriptomic findings and aligns with prior work showing that inflammatory activation within neural stem and progenitor cells can suppress neurogenesis and lead to a decline in DCX-positive neurons (Shariq et al., 2021; Sierra et al., 2015). Treatment with GSM-15606 under DEP exposure significantly increased DCX-positive cell numbers relative to DEP alone, indicating a partial recovery rather than complete restoration of neurogenesis (Fig 4B).

**Figure 4.**
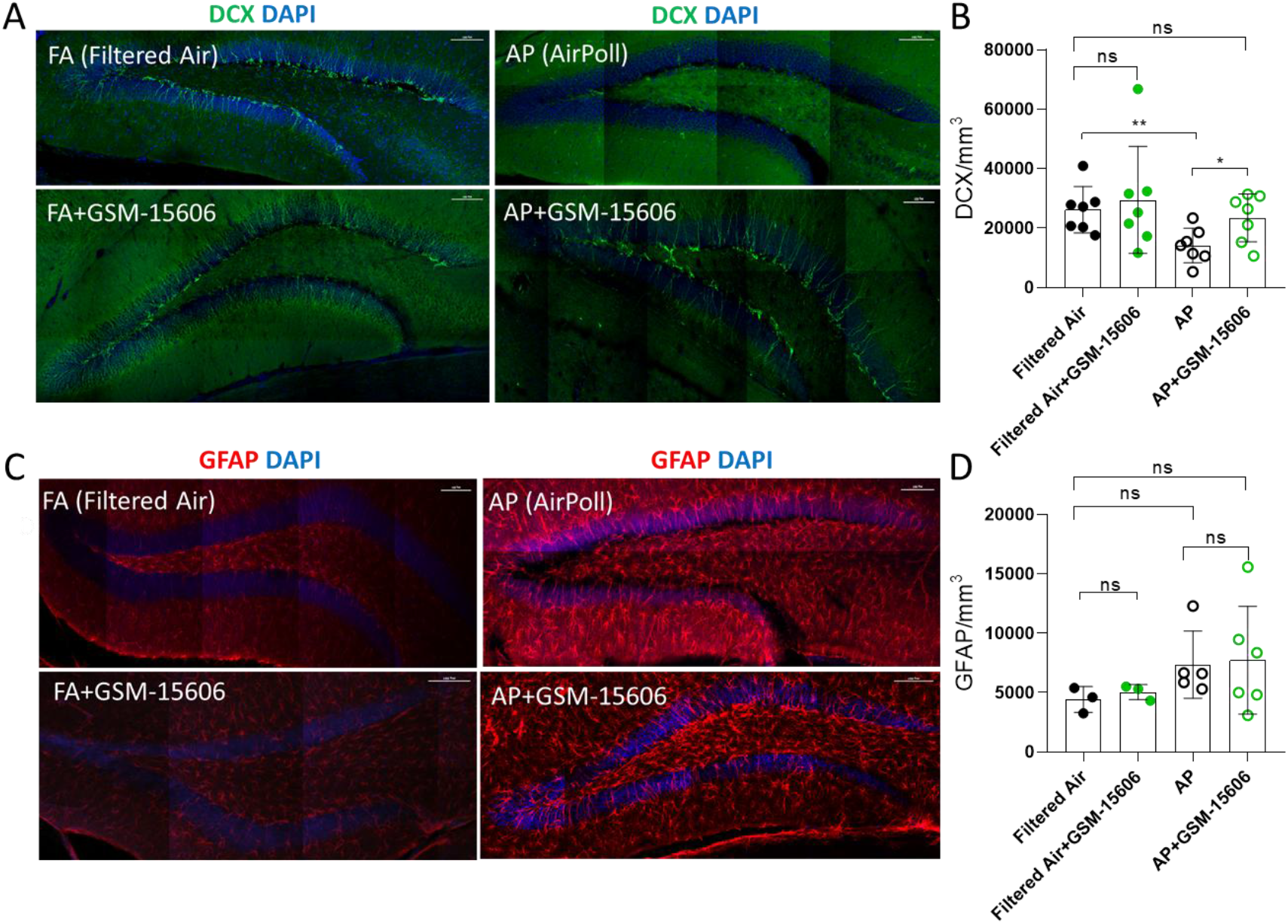
Diesel exhaust particle exposure suppresses hippocampal neurogenesis with partial recovery by GSM-15606. (A)Representative immunofluorescence images of DCX-positive immature neurons in the dentate gyrus across experimental groups: filtered air (FA), FA + GSM-15606, AirPoll (AP), and AP + GSM-15606. (B) Quantification of DCX-positive cell density (cells/mm^3^) shows a significant reduction following AP exposure compared with FA controls, with a significant increase in DCX-positive cells in the AP + GSM-15606 group relative to AP alone, indicating partial recovery of neurogenesis. (C) Representative immunofluorescence images of GFAP-positive astrocytes in the dentate gyrus across the same experimental groups. (D) Quantification of GFAP-positive astrocyte density (cells/mm^3^) reveals no statistically significant differences between groups, although a consistent upward trend is observed in AP-exposed animals compared with FA controls. GSM-15606 treatment does not alter GFAP-positive cell numbers under either FA or AP conditions. Data are presented as individual data points with mean ± SD. Statistical significance is indicated where applicable; ns denotes not significant. each dot represents an animal, (unpaired *t* test, \**P* < 0.01, \*\**P* < 0.001). Scale bars, 100μm.

In parallel, staining for GFAP-positive astrocytes did not reveal statistically significant differences across experimental groups (Fig 4C-D). Nevertheless, a consistent upward trend in GFAP-positive cell density was observed in DEP-exposed animals compared with filtered air controls, suggesting a tendency toward increased astrocytic representation within the neurogenic niche. GFAP levels in both GSM-15606-treated groups remained comparable to filtered air, indicating stability of astrocyte numbers within the sensitivity of this analysis (Fig 4D).

Together, these findings confirm at the tissue level that DEP exposure suppresses newborn neuron production and is accompanied by a trend toward astrocytic expansion in the hippocampus. The partial recovery of DCX-positive neurons with GSM-15606 treatment supports a selective mitigation of pollution-induced neurogenic impairment, consistent with its modulatory effects on inflammatory signaling identified at the single-cell level.

### 4.5. GSM-15606 selectively reduces glial inflammation but does not alter the neural stem cell response

To test whether GSM-15606 modifies pollution-driven transcriptional changes, we quantified cell type-specific gene set scores and then examined gene-level rescue using the pollution index and rescue index framework. In microglia, the AirPoll inflammation gene set score increased with DEP exposure and was reduced with GSM-15606 treatment, with AP + GSM shifting toward the filtered air distribution (Fig 5A). Consistent with this global shift, the rescue scatter plot identifies a focused subset of pollution-responsive microglial transcripts that meet criteria for significant rescue. The rescued genes highlighted in Fig. 5D include mt-Cytb, Extl3, Chd9, Adam10, Tmem119, Cep170, Cmtm6, Adgrg1, Itgam, Prpf40a, F11r, Rsrp1, and Lpcat2.

**Figure 5.**
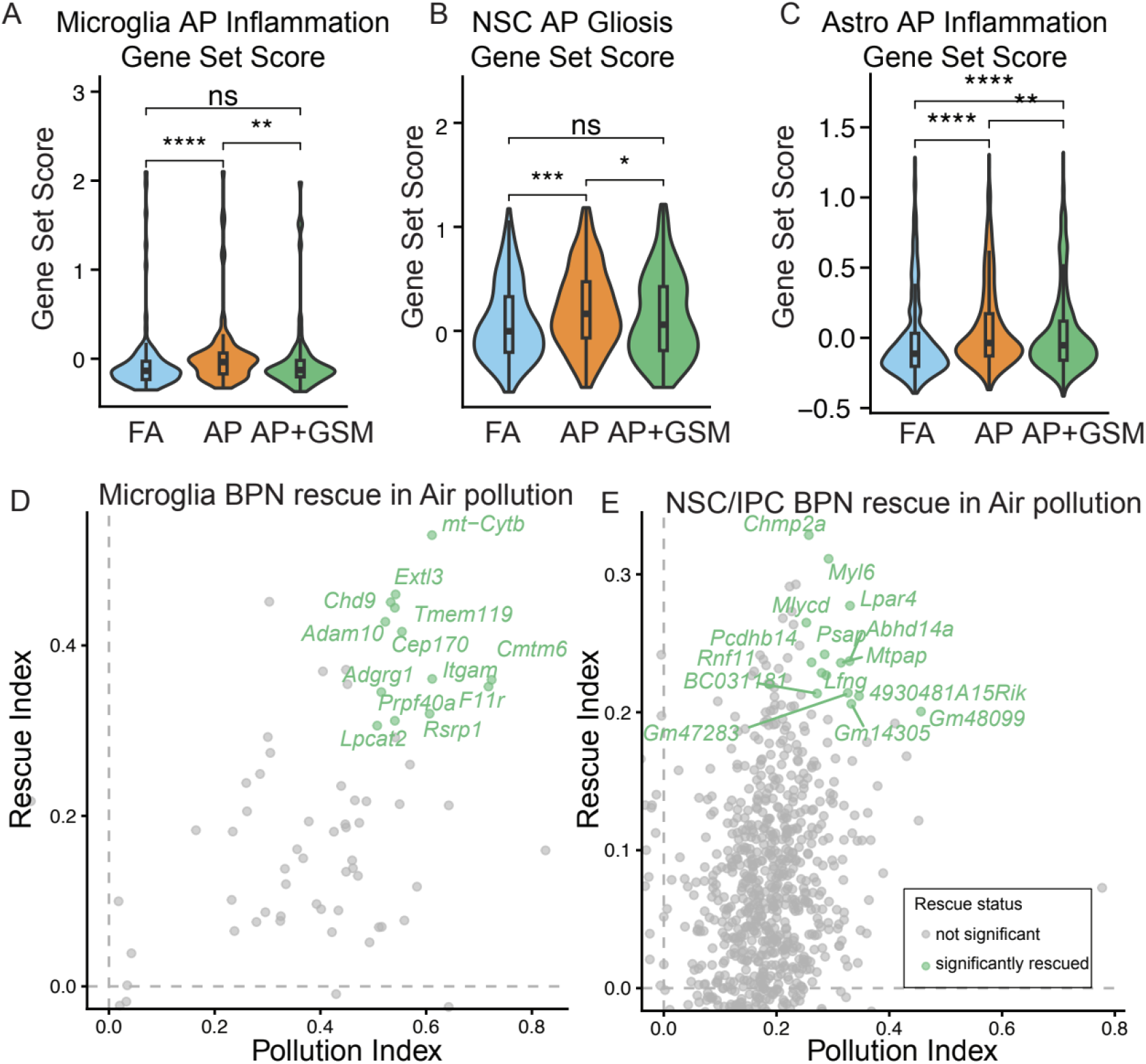
GSM-15606 attenuates air-pollution–associated inflammatory phenotypes. (A) Microglia inflammation gene set scores across conditions, showing reduced inflammatory activation with GSM-15606 in the air pollution context. Pairwise statistical testing between the indicated condition groups was performed using two-sided Wilcoxon rank-sum tests and significance is displayed on the plot as p.signif (ns, *, **, ***, ****). (B) Microglia log fold-change (LFC) “rescue quadrant” plot comparing air pollution vs Filtered Air (x-axis) against air pollution + GSM-15606 vs air pollution (y-axis), illustrating that air pollution-upregulated inflammatory genes tend to shift in the opposite direction with GSM-15606 treatment. (C) Astrocyte module scores for an air pollution-upregulated inflammatory gene set (left) and a GSM-15606-down-regulated inflammatory gene set (right), suggesting extremely limited overlap but distinct, significant rescue-associated gene subsets. (D) Astrocyte LFC–LFC quadrant plot showing the overall directional relationship between air pollution effects and air pollution + GSM-15606 effects across gene subsets.

Several of these genes map to microglial identity and immune interface functions, including Tmem119 as a microglial lineage marker, Itgam as an innate immune receptor component, and Lpcat2 as a lipid remodeling enzyme linked to inflammatory activation states. This pattern supports partial normalization of microglial perturbation rather than a uniform reversal of every inflammation-linked transcript.

In NSC and intermediate progenitor cells, DEP increased the gliosis-associated gene set score, and GSM-15606 did not produce a comparable normalization at the gene set level (Fig 5B). Nonetheless, the gene-level rescue plot in Fig. 5E indicates that GSM-15606 significantly rescues a discrete group of NSC and IPC transcripts, including Chmp2a, Myl6, Mlycd, Lpar4, Abhd14a, Psap, Mtpap, Lfng, Pcdhb14, Rnf11, BC031181, Gm47283, Gm14305, 4930481A15Rik, and Gm48099.

Next, we examined the astrocyte transcriptional response to diesel exhaust particle exposure and assessed the extent to which GSM-15606 modulates these changes. Consistent with the reduction in astrocyte inflammatory burden, analysis of GSM-15606 responsive transcripts revealed selective normalization of DEP-induced astrocyte activation programs (Fig S4). Several genes that were upregulated by DEP and are associated with astrocyte reactivity and inflammatory signaling were downregulated following GSM-15606 treatment (Fig 5C). These included Ackr3 (CXCR7), which regulates chemokine responsiveness and astrocyte migration in inflammatory and injury contexts (Li et al., 2019; van der Meer et al., 2010), and Gsn (gelsolin), a mediator of cytoskeletal remodeling that is increased in reactive astrocytes and linked to inflammatory motility and structural plasticity (Endres et al., 1998; Kamada et al., 2017). Additional stress- and inflammation-associated genes, such as Birc3, an NF-κB-regulated survival factor (Vallabhapurapu & Karin, 2009), Otulin, a regulator of ubiquitin-dependent inflammatory signaling (Keusekotten et al., 2013), Gpnmb, which marks injury-responsive astrocytes and microglia (Neal et al., 2018), and A2m, an acute-phase protein upregulated during neuroinflammation (Wyss-Coray et al., 2003), were also reduced. In contrast, canonical markers of strong astrocyte activation, including C3, immediate early genes (Fos, Fosb, Junb, Jund), and the inflammatory transcription factor Cebpb, were not substantially altered, indicating that GSM-15606 dampens specific inflammatory signaling pathways without fully reverting astrocytes to a quiescent state (Liddelow et al., 2017; Scieszka et al., 2022).

The identity of these rescued genes is notable because they largely cluster around cellular homeostasis and intracellular handling pathways, such as endosomal and lysosomal trafficking (Chmp2a, Psap) and mitochondrial or metabolic regulation (Mtpap, Mlycd), rather than the core fate-biasing inflammatory axis emphasized earlier for NSCs (Boda et al., 2020; Bonni et al., 1997; Choi & Park, 2023; Jin et al., 2025; Nakashima et al., 1999). Importantly, the NSC transcripts we highlighted as gliosis promoting in response to DEP, including Il6st, Stat3, and Txnip, are not among the significantly rescued genes annotated in Fig. 5E, consistent with limited treatment impact on the DEP induced NSC inflammatory and fate-biasing program.

Taken together, results shows that GSM-15606 shifts pollution-associated transcription in glia toward the filtered air state and rescues a defined subset of pollution-responsive genes in microglia, NSC/IPC, and astrocyte while the NSC gliosis-associated program remains comparatively resistant at the pathway level

### 4.6. GSM-15606 shows limited immunomodulatory effects in microglia under filtered air exposure

To assess whether GSM-15606 alters baseline inflammatory states independent of pollution exposure, we examined its effects in microglia from animals maintained under filtered air conditions. GSM-15606 did not significantly change the AirPoll –associated inflammatory gene set score in microglia when compared with filtered air controls, indicating that treatment does not induce a broad pro-inflammatory transcriptional shift at baseline. This outcome suggests that GSM-15606 does not intrinsically activate microglial inflammatory programs in the absence of environmental insult.

At the gene level, differential expression analysis revealed a limited number of transcripts that were modestly altered by GSM-15606 under filtered air conditions. These included immune- and stress-related genes such as *Slc40a1, Cfh, Ptgs2, Cd36, Cd38, Cxcl2, P2rx7, Zfp36, Ccl7*, and *Hspa1a/b* (Fig 6A). Several of these genes are involved in iron handling, complement regulation, lipid sensing, purinergic signaling, and immediate-early stress responses (Fig 6B), pathways that are known to fluctuate with immune tone rather than reflecting sustained inflammatory activation (Costa et al., 2017; Cory-Slechta & Sobolewski, 2023). Importantly, the magnitude and scope of these changes were restricted, and canonical markers of chronic microglial activation and antigen presentation were not broadly induced.

**Figure 6.**
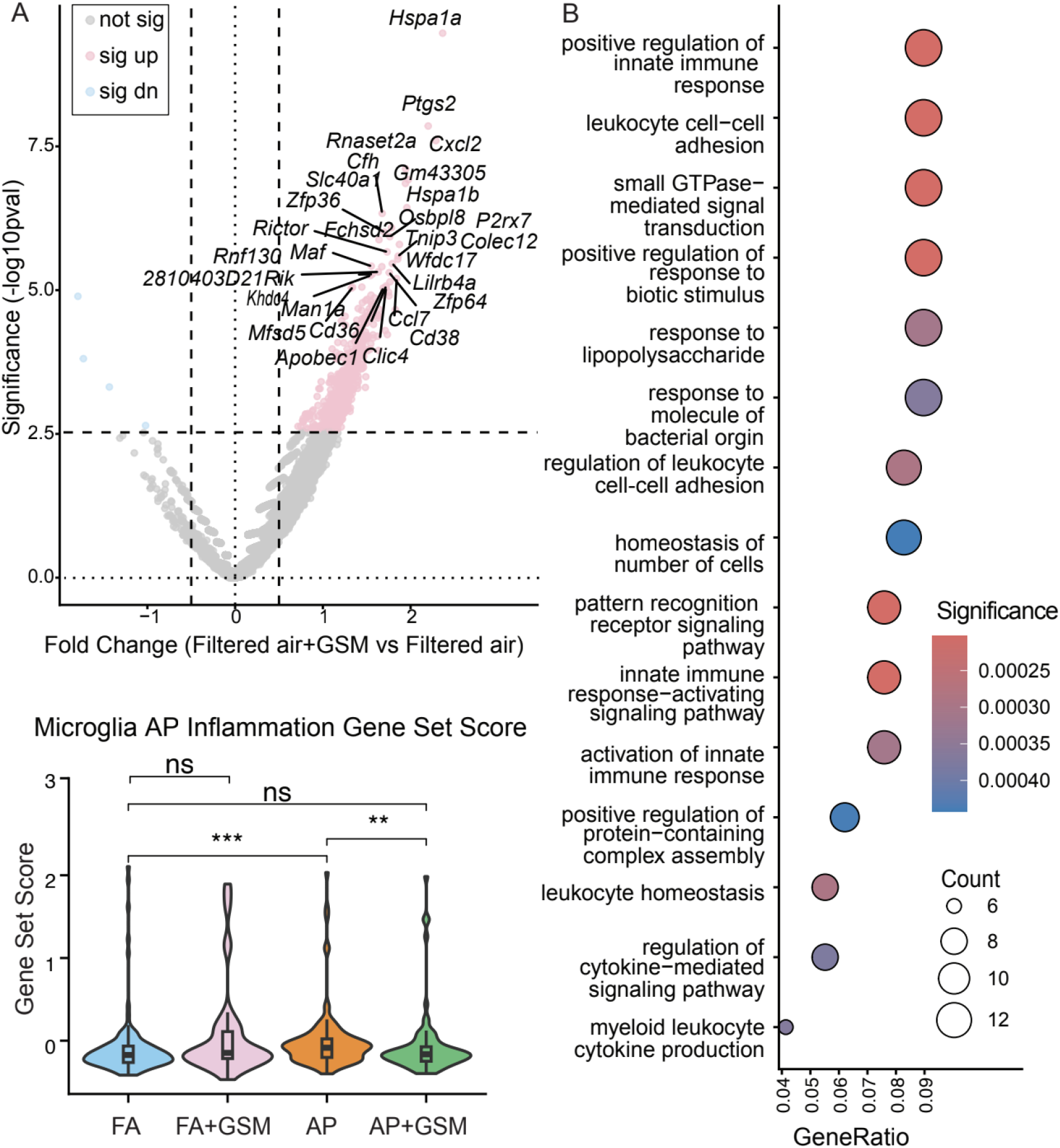
GSM-15606 does not induce broad microglial inflammatory activation under filtered air conditions. (A) Differential gene expression analysis of microglia from animals maintained under filtered air conditions comparing vehicle and GSM-15606 treatment. (B) Gene ontology enrichment analysis of GSM-15606-responsive microglial transcripts under filtered air conditions reveals enrichment of immune-adjacent processes. Across panels, GSM-15606 treatment does not significantly alter the air pollution associated inflammatory gene set score in microglia under filtered air conditions

Gene ontology enrichment analysis of GSM-responsive transcripts under filtered air conditions identified immune-adjacent processes, including pattern recognition receptor signaling, regulation of innate immune responses, and leukocyte cell-cell adhesion (Fig 6B). However, these enrichments arose from a small set of differentially expressed genes rather than a global transcriptional reprogramming, supporting the interpretation that GSM-15606 exerts modest immunomodulatory effects without driving pathological microglial activation. Together, these findings indicate that GSM-15606 does not elicit a pronounced inflammatory response in microglia under baseline conditions, supporting its use as a modulatory intervention with minimal off-target inflammatory consequences in the absence of AirPoll exposure.

## 5. Discussion

AirPoll is increasingly recognized as a major environmental determinant of brain health, with converging epidemiological and experimental evidence linking long-term exposure to particulate matter and traffic-related pollutants to accelerated cognitive decline and increased dementia risk (Cory-Slechta & Sobolewski, 2023; Mork et al., 2023; Shi et al., 2020; Wilker et al., 2023). Emerging mechanistic work suggests that these effects arise from cumulative, mixture-based exposures that engage shared pathological pathways, including neuroinflammation, oxidative stress, blood–brain barrier dysfunction, and activation of microglia and astrocytes—processes that overlap with early features of Alzheimer’s disease (Calderón-Garcidueñas et al., 2008; Costa et al., 2017; Iaccarino, La Joie, Edwards, et al., 2021). However, the cell-type specificity of these responses within the hippocampal neurogenic niche, and the extent to which they can be selectively modulated by therapeutic intervention, remain incompletely defined.

Here, we used single-cell transcriptomics coupled with histological validation to resolve how diesel exhaust particle exposure reshapes the hippocampal neurogenic niche. This approach revealed pronounced mosaicism in pollution-induced responses, uncovering distinct transcriptional programs across microglia, astrocytes, and neural stem cells that are not apparent in bulk analyses. Our findings show that AirPoll activates coordinated but cell-type-specific inflammatory states that converge on impaired neurogenesis and increased gliosis.

Microglia exhibited robust activation of inflammatory and antigen-presenting programs, including cytokine and chemokine signaling, stress-responsive transcriptional regulators, and induction of MHC class II components. These changes align with prior work demonstrating that diesel and particulate exposures promote sustained microglial activation and inflammatory remodeling (Levesque et al., 2011; Sahu et al., 2021; Zhang et al., 2023). Astrocytes also exhibited a shift toward a reactive transcriptional state following DEP exposure, characterized by increased gene expression of cytokine and chemokine responsiveness, cytoskeletal remodeling, and stress-associated signaling rather than overt complement-driven activation. While classical markers of strong astrocyte reactivity were not uniformly elevated, the induction of inflammatory response regulators and immediate early genes indicates engagement of an environmentally responsive astrocyte program. These transcriptional changes were accompanied by a trend toward increased GFAP-positive astrocytes at the tissue level, although astrocyte numbers did not differ significantly across conditions. Together, these findings suggest that air pollution elicits a moderate but coordinated astrocytic inflammatory response without necessarily driving robust cellular expansion. In combination with microglial activation and inflammatory signaling, these astrocyte responses contribute to a hippocampal environment shaped by inflammatory stress and diminished support for neurogenesis (Kilian et al., 2023; Scieszka et al., 2022).

Importantly, neural stem cells also adapted a pollution-induced inflammatory and stress-responsive identity. Upregulation of IL6ST and STAT3 implicates activation of a pathway known to bias neural precursor fate decisions and to support gliotic responses during injury and inflammation (Nakashima et al., 1999; O’Callaghan et al., 2014; Taga & Fukuda, 2005). Increased TXNIP further supports an oxidative and inflammatory state within the NSC compartment, consistent with mechanisms that suppress adult neurogenesis (E. H. Choi & Park, 2023; Zhou et al., 2010). These NSC-intrinsic signatures align with the concept that stem and progenitor populations can harbor latent inflammatory potential that restricts neurogenesis under pathological conditions (Shariq et al., 2021). Consistent with these transcriptional shifts, immunohistochemical analysis revealed a significant reduction in DCX-positive immature neurons within the dentate gyrus following DEP exposure, confirming that AirPoll suppresses neuronal output at the tissue level. This loss of newborn neurons aligns with prior evidence linking inflammatory activation within the neurogenic niche to impaired neurogenesis (Costa et al., 2017; Kilian et al., 2023; Sahu et al., 2021; Shariq et al., 2021). In contrast, GFAP-positive astrocyte numbers did not show statistically significant differences across experimental groups. Nevertheless, a consistent trend toward increased GFAP-positive cells was observed in DEP-exposed animals, suggesting a shift toward astrocytic reactivity that parallels the inflammation-associated transcriptional signatures detected at single-cell resolution. Together, these findings indicate that DEP exposure robustly impairs neurogenesis while eliciting more subtle, transcriptionally evident changes in astrocyte state that may precede overt gliosis.

Treatment with the gamma secretase modulator GSM-15606 reversed a substantial portion of the DEP-induced inflammatory program in microglia and improved astrocyte states at both transcriptional and cellular levels. This phenotype is consistent with prior reports that GSM-15606 and related gamma secretase modulators reduce glial reactivity and improve pathological outcomes in Alzheimer’s disease models without broadly disrupting essential signaling (Prikhodko et al., 2020; Rynearson et al., 2021; Wagner et al., 2017). In the context of pollution exposure, our data suggest that GSM-15606 can dampen inflammatory activation in key glial populations that shape the neurogenic niche and contribute to synaptic and circuit vulnerability. Importantly, analysis of GSM-15606 treatment under filtered air conditions revealed no evidence of pathological microglial activation. Although gene ontology analysis identified enrichment for innate immune and LPS-response terms, these changes were limited in magnitude and did not include canonical markers of inflammatory microglial activation, such as pro-inflammatory cytokines and chemokines (Il1b, Tnf, Il6, Cxcl2) or antigen-presentation and MHC class II genes (H2-Aa, H2-Ab1, H2-Eb1, Cd74). The absence of these established activation markers, together with the lack of gliosis compared with DEP-exposed animals, supports the interpretation that GSM-15606 exerts modest immunomodulatory effects rather than inducing aggressive inflammatory activation, reinforcing its suitability as a modulatory intervention with minimal off-target inflammatory consequences in the absence of AirPoll exposure.

In contrast, the NSC transcriptional response to pollution was not rescued by GSM-15606. Core pollution-induced inflammatory and oxidative stress transcripts within NSCs remained elevated despite treatment, indicating that NSCs sustain an intrinsic stress-responsive state that is comparatively resistant to this intervention. This divergence between glial rescue and NSC persistence has important implications. It suggests that reducing glial activation alone may be insufficient to fully restore neurogenic output if NSCs continue to maintain inflammatory fate-biasing programs. More broadly, it points to a two-compartment model in which glia act as highly responsive amplifiers of environmental inflammation, while NSCs may encode longer-lived inflammatory memory that continues to bias the niche toward gliosis and reduced neuronal production.

The broader relevance of these findings extends to cognitive aging and Alzheimer’s disease. Adult hippocampal neurogenesis contributes to learning, memory, and cognitive flexibility, and its disruption is increasingly linked to early cognitive vulnerability (Aimone et al., 2011; Anacker et al., 2018; S. H. Choi et al., 2018; Creer et al., 2010; Frechou et al., 2024; Moreno-Jiménez et al., 2019; Shariq et al., 2021; Tobin et al., 2019; Van Praag et al., 1999, Ammothumkandy et al., 2024). Many of the pathways activated by AirPoll inthis study overlap with those implicated in Alzheimer’s pathogenesis, including microglial inflammatory reprogramming, complement activation, oxidative stress, and astrocyte reactivity (Cacciottolo et al., 2017; Calderón-Garcidueñas et al., 2008; Iaccarino, La Joie, Edwards, et al., 2021). By simultaneously suppressing neurogenesis and reinforcing gliosis, AirPoll may lower resilience of hippocampal circuits, thereby amplifying susceptibility to dementia-related processes. Our findings further suggest that therapeutics capable of dampening glial activation may reduce a key component of this vulnerability, while additional strategies may be required to directly reprogram NSC-intrinsic inflammatory pathways.

In summary, our study provides a single-cell framework for understanding how AirPoll disrupts the hippocampal neurogenic niche through distinct yet convergent inflammatory programs across microglia, astrocytes, and neural stem cells. By integrating transcriptomic and cellular readouts, we demonstrate that GSM-15606 selectively attenuates AirPoll -induced glial activation and rescues neurogenesis (Fig. 4B), while NSC inflammatory programs persist. These results position AirPoll as a modifiable risk factor for brain aging and dementia and highlight the need for cell-type-specific strategies (Li & others, 2025) that address both glial inflammation and NSC-intrinsic inflammatory states to preserve hippocampal plasticity.

## Funding

This work was supported by NIH (R01AG076956 to M.A.B; RF1NS130681 to W.J.M and C.S, F31AG084302 to J.N.L., T32HD060549 training grant to W.P., J.N.L, A.I., W.N.), Cure Alzheimer’s Fund to W.N., C.E.F, and M.A.B, Simons Strauss Foundation to M.A.B., AFAR Research Scholarship for Research in Biology of Aging to J.N.L, A.I, and W.N., California Institute for Regenerative Medicine (EDUC4 Postdoctoral training grant) to M.S. M.B. was partially supported by the USC Provost’s Research Enhancement Fellowship.

## Author contributions

M.S., C.E.B., M.A.T., and M.A.B conceived the project. X.C., J.G-L., W.J.M., C.E.B., M.A.T., and M.A.B designed the experiments. W.P., X.C., W.X, J.G-L, L.P, J.N.L, A.I., K.S., W.N performed the experiments. M.S., W.P., X.C., M.A.B analyzed and compiled the data. M.S., W.P., X.C, C.E.B., M.A.T., and M.A.B wrote the manuscript. C.E.B., M.A.T., and M.A.B supervised the project.

## Notes

### Competing Interest Statement

The authors have declared no competing interest.

## References

Aimone, J. B., Deng, W., & Gage, F. H. (2011). Resolving New Memories: A Critical Look at the Dentate Gyrus, Adult Neurogenesis, and Pattern Separation. In Neuron (Vol. 70, Issue 4). 10.1016/j.neuron.2011.05.010

Ammothumkandy, A., Corona, L., Ravina, K., Wolseley, V., Nelson, J., Atai, N., Abedi, A., Jimenez, N., Armacost, M., D’Orazio, L. M., Zuverza-Chavarria, V., Cayce, A., McCleary, C., Nune, G., Kalayjian, L., Lee, D. J., Lee, B., Chow, R. H., Heck, C., … Bonaguidi, M. A. (2024). Human adult neurogenesis loss corresponds with cognitive decline during epilepsy progression. Cell Stem Cell. 10.1016/J.STEM.2024.11.002

Anacker, C., Luna, V. M., Stevens, G. S., Millette, A., Shores, R., Jimenez, J. C., Chen, B., & Hen, R. (2018). Hippocampal neurogenesis confers stress resilience by inhibiting the ventral dentate gyrus. Nature, 559(7712), 98–102. 10.1038/s41586-018-0262-4

Bennett, M. L., Bennett, F. C., Liddelow, S. A., Ajami, B., Zamanian, J. L., Fernhoff, N. B., Mulinyawe, S. B., Bohlen, C. J., Adil, A., Tucker, A., Weissman, I. L., Chang, E. F., Li, G., Grant, G. A., Hayden Gephart, M. G., & Barres, B. A. (2016). New tools for studying microglia in the mouse and human CNS. Proceedings of the National Academy of Sciences of the United States of America, 113(12). 10.1073/pnas.1525528113

Block, M. L., & Kodavanti, U. P. (2021). The Use of Standardized Diesel Exhaust Particles in Alzheimer’s Disease Research. In Journal of Alzheimer’s Disease (Vol. 84, Issue 2). 10.3233/JAD-215201

Boda, E., Rigamonti, A. E., & Bollati, V. (2020). Understanding the effects of air pollution on neurogenesis and gliogenesis in the growing and adult brain. In Current Opinion in Pharmacology (Vol. 50). 10.1016/j.coph.2019.12.003

Bonni, A., Sun, Y., Nadal-Vicens, M., Bhatt, A., Frank, D. A., Rozovsky, I., Stahl, N., Yancopoulos, G. D., & Greenberg, M. E. (1997). Regulation of gliogenesis in the central nervous system by the JAK-STAT signaling pathway. Science, 278(5337). 10.1126/science.278.5337.477

Cacciottolo, M., Wang, X., Driscoll, I., Woodward, N., Saffari, A., Reyes, J., Serre, M. L., Vizuete, W., Sioutas, C., Morgan, T. E., Gatz, M., Chui, H. C., Shumaker, S. A., Resnick, S. M., Espeland, M. A., Finch, C. E., & Chen, J. C. (2017). Particulate air pollutants, APOE alleles and their contributions to cognitive impairment in older women and to amyloidogenesis in experimental models. Translational Psychiatry, 7(1). 10.1038/tp.2016.280

Calderón-Garcidueñas, L., Solt, A. C., Henríquez-Roldán, C., Torres-Jardón, R., Nuse, B., Herritt, L., Villarreal-Calderón, R., Osnaya, N., Stone, I., García, R., Brooks, D. M., González-Maciel, A., Reynoso-Robles, R., Delgado-Chávez, R., & Reed, W. (2008). Long-term air pollution exposure is associated with neuroinflammation, an altered innate immune response, disruption of the blood-brain barrier, ultrafine particulate deposition, and accumulation of amyloid β-42 and α-synuclein in children and young adults. Toxicologic Pathology, 36(2). 10.1177/0192623307313011

Cheng, L., Lau, W. K. W., Fung, T. K. H., Lau, B. W. M., Chau, B. K. H., Liang, Y., Wang, Z., So, K. F., Wang, T., Chan, C. C. H., & Lee, T. M. C. (2017). PM2.5 Exposure Suppresses Dendritic Maturation in Subgranular Zone in Aged Rats. Neurotoxicity Research, 32(1). 10.1007/s12640-017-9710-4

Choi, E. H., & Park, S. J. (2023). TXNIP: A key protein in the cellular stress response pathway and a potential therapeutic target. In Experimental and Molecular Medicine (Vol. 55, Issue 7). 10.1038/s12276-023-01019-8

Choi, S. H., Bylykbashi, E., Chatila, Z. K., Lee, S. W., Pulli, B., Clemenson, G. D., Kim, E., Rompala,, Oram, M. K., Asselin, C., Aronson, J., Zhang, C., Miller, S. J., Lesinski, A., Chen, J. W., Kim, D. Y., Praag, H. Van, Spiegelman, B. M., Gage, F. H., & Tanzi, R. E. (2018). Combined adult neurogenesis and BDNF mimic exercise effects on cognition in an Alzheimer’s mouse model. Science (New York, N.Y.), 361(6406). 10.1126/SCIENCE.AAN8821

Cory-Slechta, D. A., & Sobolewski, M. (2023). Neurotoxic effects of air pollution: an urgent public health concern. In Nature Reviews Neuroscience (Vol. 24, Issue 3). 10.1038/s41583-022-00672-8

Costa, L. G., Cole, T. B., Coburn, J., Chang, Y. C., Dao, K., & Roqué, P. J. (2017). Neurotoxicity of traffic-related air pollution. NeuroToxicology, 59. 10.1016/j.neuro.2015.11.008

Creer, D. J., Romberg, C., Saksida, L. M., Van Praag, H., & Bussey, T. J. (2010). Running enhances spatial pattern separation in mice. Proceedings of the National Academy of Sciences of the United States of America, 107(5). 10.1073/pnas.0911725107

Farahani, V. J., Pirhadi, M., & Sioutas, C. (2021). Are standardized diesel exhaust particles (DEP) representative of ambient particles in air pollution toxicological studies? Science of the Total Environment, 788. 10.1016/j.scitotenv.2021.147854

Finch, C. E., & Thorwald, M. A. (2024). Inhaled Pollutants of the Gero-Exposome and Later-Life Health. In Journals of Gerontology - Series A Biological Sciences and Medical Sciences (Vol. 79, Issue 7). 10.1093/gerona/glae107

Frechou, M. A., Martin, S. S., McDermott, K. D., Huaman, E. A., Gökhan, Ş., Tomé, W. A., Coen-Cagli, R., & Gonçalves, J. T. (2024). Adult neurogenesis improves spatial information encoding in the mouse hippocampus. Nature Communications, 15(1). 10.1038/s41467-024-50699-x

Godoy-Lugo, J. A., Hicks, D., Durra, S., Massey, E. R., Shkirkova, K., Sauri, A. I., Kerstiens, E., Chen, S., Zhao, L., Sioutas, C., Mack, W. J., Finch, C. E., & Thorwald, M. A. (2025). Air pollution decreases postsynaptic PSD-95 and NMDA receptor subunits in synaptosomes from mouse cerebral cortex. Environmental Pollution, 383. 10.1016/j.envpol.2025.126845

Godoy-Lugo, J. A., Thorwald, M. A., Cacciottolo, M., D’Agostino, C., Chakhoyan, A., Sioutas, C., Tanzi, R. E., Rynearson, K. D., & Finch, C. E. (2024). Air pollution amyloidogenesis is attenuated by the gamma-secretase modulator GSM-15606. Alzheimer’s and Dementia, 20(9). 10.1002/alz.14086

Hart J, Harris L. Protocol to analyze changes in hippocampal neural stem cell quiescence from single-cell RNA sequencing data. STAR Protoc. 2025 Nov 24;6(4):104226. doi: 10.1016/j.xpro.2025.104226. Epub ahead of print. PMID: 41289073; PMCID: PMC12682022.

Hartz, A. M. S., Bauer, B., Block, M. L., Hong, J.-S., & Miller, D. S. (2008). Diesel exhaust particles induce oxidative stress, proinflammatory signaling, and P-glycoprotein up-regulation at the blood-brain barrier. The FASEB Journal, 22(8). 10.1096/fj.08-106997

Iaccarino, L., La Joie, R., Edwards, L., Strom, A., Schonhaut, D. R., Ossenkoppele, R., Pham, J., Mellinger, T., Janabi, M., Baker, S. L., Soleimani-Meigooni, D., Rosen, H. J., Miller, B. L., Jagust, W. J., & Rabinovici, G. D. (2021). Spatial Relationships between Molecular Pathology and Neurodegeneration in the Alzheimer’s Disease Continuum. Cerebral Cortex, 31(1). 10.1093/cercor/bhaa184

Iaccarino, L., La Joie, R., Lesman-Segev, O. H., Lee, E., Hanna, L., Allen, I. E., Hillner, B. E., Siegel, A., Whitmer, R. A., Carrillo, M. C., Gatsonis, C., & Rabinovici, G. D. (2021). Association between Ambient Air Pollution and Amyloid Positron Emission Tomography Positivity in Older Adults with Cognitive Impairment. JAMA Neurology, 78(2). 10.1001/jamaneurol.2020.3962

Jaiswal, C., & Singh, A. K. (2024). Particulate matter exposure and its consequences on hippocampal neurogenesis and cognitive function in experimental models. In Environmental Pollution (Vol. 363). 10.1016/j.envpol.2024.125275

Jin, N., Lee, J., Park, S. Y., & Han, J. S. (2025). NOTCH1-STAT3 signaling axis regulates astrocytic differentiation of hippocampal neural stem/progenitor cells. Biochemical and Biophysical Research Communications, 765. 10.1016/j.bbrc.2025.151844

Kilian, J. G., Mejias-Ortega, M., Hsu, H. W., Herman, D. A., Vidal, J., Arechavala, R. J., Renusch, S., Dalal, H., Hasen, I., Ting, A., Rodriguez-Ortiz, C. J., Lim, S. L., Lin, X., Vu, J., Saito, T., Saido, T. C., Kleinman, M. T., & Kitazawa, M. (2023). Exposure to quasi-ultrafine particulate matter accelerates memory impairment and Alzheimer’s disease-like neuropathology in the AppNL-G-F knock-in mouse model. Toxicological Sciences, 193(2). 10.1093/toxsci/kfad036

Levesque, S., Surace, M. J., McDonald, J., & Block, M. L. (2011). Air pollution and the brain: Subchronic diesel exhaust exposure causes neuroinflammation and elevates early markers of neurodegenerative disease. Journal of Neuroinflammation, 8. 10.1186/1742-2094-8-105

Li, Y., Serras, C. P., Blumenfeld, J., Xie, M., Hao, Y., Deng, E., … & Sirota, M. (2025). Cell-type-directed network-correcting combination therapy for Alzheimer’s disease. Cell, 188(20), 5516–5534.

Liang Z, Jin N, Guo W. Neural stem cell heterogeneity in adult hippocampus. Cell Regen. 2025 Mar 7;14(1):6. doi: 10.1186/s13619-025-00222-4. PMID: 40053275; PMCID: PMC11889326.

Moreno-Jiménez, E. P., Flor-García, M., Terreros-Roncal, J., Rábano, A., Cafini, F., Pallas-Bazarra, N., Ávila, J., & Llorens-Martín, M. (2019). Adult hippocampal neurogenesis is abundant in neurologically healthy subjects and drops sharply in patients with Alzheimer’s disease. Nature Medicine, 25(4), 554–560. 10.1038/s41591-019-0375-9

Morgan, T. E., Davis, D. A., Iwata, N., Tanner, J. A., Snyder, D., Ning, Z., Kam, W., Hsu, Y. T., Winkler, J. W., Chen, J. C., Petasis, N. A., Baudry, M., Sioutas, C., & Finch, C. E. (2011). Glutamatergic neurons in rodent models respond to nanoscale particulate urban air pollutants in vivo and in vitro. Environmental Health Perspectives, 119(7). 10.1289/ehp.1002973

Morimoto, R., Shindou, H., Oda, Y., & Shimizu, T. (2010). Phosphorylation of lysophosphatidylcholine acyltransferase 2 at Ser 34 enhances platelet-activating factor production in endotoxin-stimulated macrophages. Journal of Biological Chemistry, 285(39). 10.1074/jbc.M110.147025

Mork, D., Braun, D., & Zanobetti, A. (2023). Time-lagged relationships between a decade of air pollution exposure and first hospitalization with Alzheimer’s disease and related dementias. Environment International, 171. 10.1016/j.envint.2022.107694

Nakashima, K., Yanagisawa, M., Arakawa, H., Kimura, N., Hisatsune, T., Kawabata, M., Miyazono, K., & Taga, T. (1999). Synergistic signaling in fetal brain by STAT3-Smad1 complex bridged by p300. Science, 284(5413). 10.1126/science.284.5413.479

O’Callaghan, J. P., Kelly, K. A., VanGilder, R. L., Sofroniew, M. V., & Miller, D. B. (2014). Early activation of STAT3 regulates reactive astrogliosis induced by diverse forms of neurotoxicity. PLoS ONE, 9(7). 10.1371/journal.pone.0102003

Prikhodko, O., Rynearson, K. D., Sekhon, T., Mante, M. M., Nguyen, P. D., Rissman, R. A., Tanzi, R. E., & Wagner, S. L. (2020). The GSM BPN-15606 as a Potential Candidate for Preventative Therapy in Alzheimer’s Disease. Journal of Alzheimer’s Disease, 73(4). 10.3233/JAD-190442

Roberts, A. L., Fürnrohr, B. G., Vyse, T. J., & Rhodes, B. (2016). The complement receptor 3 (CD11b/CD18) agonist Leukadherin-1 suppresses human innate inflammatory signalling. Clinical and Experimental Immunology, 185(3). 10.1111/cei.12803

Rynearson, K. D., Ponnusamy, M., Prikhodko, O., Xie, Y., Zhang, C., Nguyen, P., Hug, B., Sawa, M., Becker, A., Spencer, B., Florio, J., Mante, M., Salehi, B., Arias, C., Galasko, D., Head, B. P., Johnson, G., Lin, J. H., Duddy, S. K., … Wagner, S. L. (2021). Preclinical validation of a potent γ-secretase modulator for Alzheimer’s disease prevention. Journal of Experimental Medicine, 218(4). 10.1084/JEM.20202560

Sahu, B., Mackos, A. R., Floden, A. M., Wold, L. E., & Combs, C. K. (2021). Particulate Matter Exposure Exacerbates Amyloid-β Plaque Deposition and Gliosis in APP/PS1 Mice. Journal of Alzheimer’s Disease, 80(2). 10.3233/JAD-200919

Satoh, J. ichi, Kino, Y., Asahina, N., Takitani, M., Miyoshi, J., Ishida, T., & Saito, Y. (2016). TMEM119 marks a subset of microglia in the human brain. Neuropathology, 36(1). 10.1111/neup.12235

Scieszka, D., Hunter, R., Begay, J., Bitsui, M., Lin, Y., Galewsky, J., Morishita, M., Klaver, Z., Wagner, J., Harkema, J. R., Herbert, G., Lucas, S., McVeigh, C., Bolt, A., Bleske, B., Canal, C. G., Mostovenko, E., Ottens, A. K., Gu, H., … Noor, S. (2022). Neuroinflammatory and Neurometabolomic Consequences from Inhaled Wildfire Smoke-Derived Particulate Matter in the Western United States. Toxicological Sciences, 186(1). 10.1093/toxsci/kfab147

Sekheri, M., Othman, A., & Filep, J. G. (2021). β2 Integrin Regulation of Neutrophil Functional Plasticity and Fate in the Resolution of Inflammation. In Frontiers in Immunology (Vol. 12). 10.3389/fimmu.2021.660760

Shariq, M., Sahasrabuddhe, V., Krishna, S., Radha, S., Nruthyathi Bellampalli, R., Dwivedi, A., Cheramangalam, R., Reizis, B., Hébert, J., & Ghosh, H. S. (2021). Adult neural stem cells have latent inflammatory potential that is kept suppressed by Tcf4 to facilitate adult neurogenesis. Science Advances, 7(21), 1–12. 10.1126/sciadv.abf5606

Shi, L., Wu, X., Danesh Yazdi, M., Braun, D., Abu Awad, Y., Wei, Y., Liu, P., Di, Q., Wang, Y., Schwartz, J., Dominici, F., Kioumourtzoglou, M. A., & Zanobetti, A. (2020). Long-term effects of PM2.5 on neurological disorders in the American Medicare population: a longitudinal cohort study. The Lancet Planetary Health, 4(12). 10.1016/S2542-5196(20)30227-8

Shkirkova, K., Demetriou, A. N., Sizdahkhani, S., Lamorie-Foote, K., Zhang, H., Morales, M., Chen, S., Zhao, L., Diaz, A., Godoy-Lugo, J. A., Zhou, B., Zhang, N., Li, A., Mack, W. J., Sioutas, C., Thorwald, M. A., Finch, C. E., Pike, C., & Mack, W. J. (2024). Microglial TLR4 Mediates White Matter Injury in a Combined Model of Diesel Exhaust Exposure and Cerebral Hypoperfusion. Stroke, 55(4). 10.1161/STROKEAHA.124.046412

Shkirkova, K., Maria, N. S. S., Anson, H., Aghaei, Y., Badami, M. M., Chakhoyan, A., Godoy-Lugo, J. A., Chung, C., Durra, S., Tang-Tan, A., Zhao, L., Demetriou, A., Morales, M., Daggupati, S., Bent, I., Park, H., Franklin, C., Chen, S., Chahine, G., … Thorwald, M. A. (2026). Induction of ferroptotic and amyloidogenic signatures linked to Alzheimers disease by chemically distinct air pollutants. BioRxiv. 10.64898/2026.01.06.696601

Sierra, A., Encinas, J. M., Deudero, J. J. P., Chancey, J. H., Enikolopov, G., Overstreet-Wadiche, L. S., Tsirka, S. E., & Maletic-Savatic, M. (2010). Microglia shape adult hippocampal neurogenesis through apoptosis-coupled phagocytosis. Cell Stem Cell. 10.1016/j.stem.2010.08.014

Taga, T., & Fukuda, S. (2005). Role of IL-6 in the neural stem cell differentiation. In Clinical Reviews in Allergy and Immunology (Vol. 28, Issue 3). 10.1385/CRIAI:28:3:249

Tobin, M. K., Musaraca, K., Disouky, A., Shetti, A., Bheri, A., Honer, W. G., Kim, N., Dawe, R. J., Bennett, D. A., Arfanakis, K., & Lazarov, O. (2019). Human Hippocampal Neurogenesis Persists in Aged Adults and Alzheimer’s Disease Patients. Cell Stem Cell, 24(6), 974-982.e3. 10.1016/j.stem.2019.05.003

Van Praag, H., Kempermann, G., & Gage, F. H. (1999). Running increases cell proliferation and neurogenesis in the adult mouse dentate gyrus. Nature Neuroscience, 2(3), 266–270. 10.1038/6368

Wagner, S. L., Rynearson, K. D., Duddy, S. K., Zhang, C., Nguyen, P. D., Becker, A., Vo, U., Masliah, D., Monte, L., Klee, J. B., Echmalian, C. M., Xia, W., Quinti, L., Johnson, G., Lin, J. H., Kim, D. Y., Mobley, W. C., Rissman, R. A., & Tanzi, R. E. (2017). Pharmacological and toxicological properties of the potent oral γ-secretase modulator BPN-15606. Journal of Pharmacology and Experimental Therapeutics, 362(1). 10.1124/jpet.117.240861

Wilker, E. H., Osman, M., & Weisskopf, M. G. (2023). Ambient air pollution and clinical dementia: systematic review and meta-analysis. BMJ, 381. 10.1136/bmj-2022-071620

Woodward, N. C., Haghani, A., Johnson, R. G., Hsu, T. M., Saffari, A., Sioutas, C., Kanoski, S. E., Finch, C. E., & Morgan, T. E. (2018). Prenatal and early life exposure to air pollution induced hippocampal vascular leakage and impaired neurogenesis in association with behavioral deficits. Translational Psychiatry, 8(1). 10.1038/s41398-018-0317-1

Yao, Y., Lv, X., Qiu, C., Li, J., Wu, X., Zhang, H., Yue, D., Liu, K., Eshak, E. S., Lorenz, T., Anstey, K. J., Livingston, G., Xue, T., Zhang, J., Wang, H., & Zeng, Y. (2022). The effect of China’s Clean Air Act on cognitive function in older adults: a population-based, quasi-experimental study. The Lancet Healthy Longevity, 3(2). 10.1016/S2666-7568(22)00004-6

Zhang, H., D’Agostino, C., Tulisiak, C., Thorwald, M. A., Bergkvist, L., Lindquist, A., Meyerdirk, L., Schulz, E., Becker, K., Steiner, J. A., Cacciottolo, M., Kwatra, M., Rey, N. L., Escobar Galvis, M. L., Ma, J., Sioutas, C., Morgan, T. E., Finch, C. E., & Brundin, P. (2023). Air pollution nanoparticle and alpha-synuclein fibrils synergistically decrease glutamate receptor A1, depending upon nPM batch activity. Heliyon, 9(4). 10.1016/j.heliyon.2023.e15622

Zhou, R., Tardivel, A., Thorens, B., Choi, I., & Tschopp, J. (2010). Thioredoxin-interacting protein links oxidative stress to inflammasome activation. Nature Immunology, 11(2). 10.1038/ni.1831

