## Supplementary data 1-4 for "Single cell profiling reveals GSM-15606 attenuates air pollution-induced inflammation and preserves hippocampal neurogenesis"

**Supplementary Information**

**Figures S1-S4.**

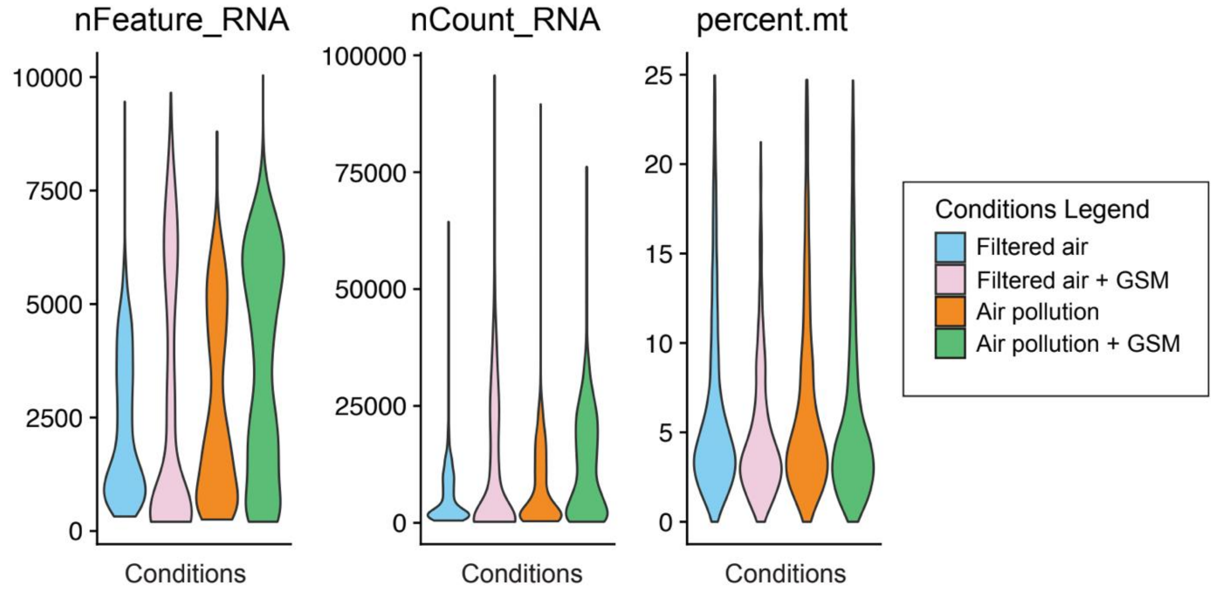


**Supplementary Figure 1: Quality control metrics for single-cell RNA-seq**
Violin plots show the distribution of number of detected genes per cell (nFeature_RNA), total UMI counts per cell (nCount_RNA), and the fraction of mitochondrial transcriptions (percent.mt). Cells with percent.mt > 25% are excluded from downstream analysis. Diesel exhaust particle (DEP) air pollution, GSM: GSM-15606 treatment.

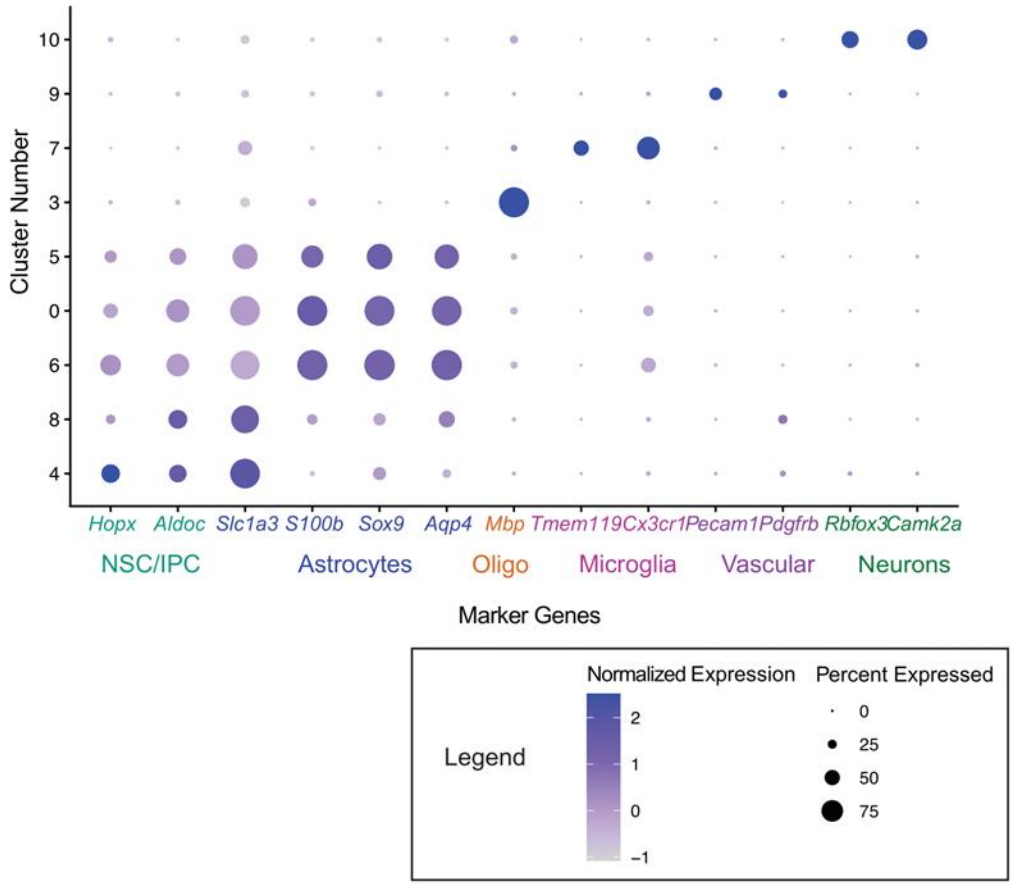


**Supplementary Figure 2: Cell type identification by canonical marker expression.** Dot plot showing scaled expression (color) and the fraction of cells expressing each marker gene (dot size) across clusters. Neural stem cells (NSCs) are identified by *Hopx* and *Aldoc*; astrocytes by *Slc1a3*, *S100b*, *Sox9*, and *Aqp4*; oligodendrocytes by *Mbp*; microglia by *Tmem119* and *Cx3cr1*; endothelial cells by *Pecam1*; pericytes by *Pdgfrb*; and neurons by *Rbfox2* and *Camk2a*.

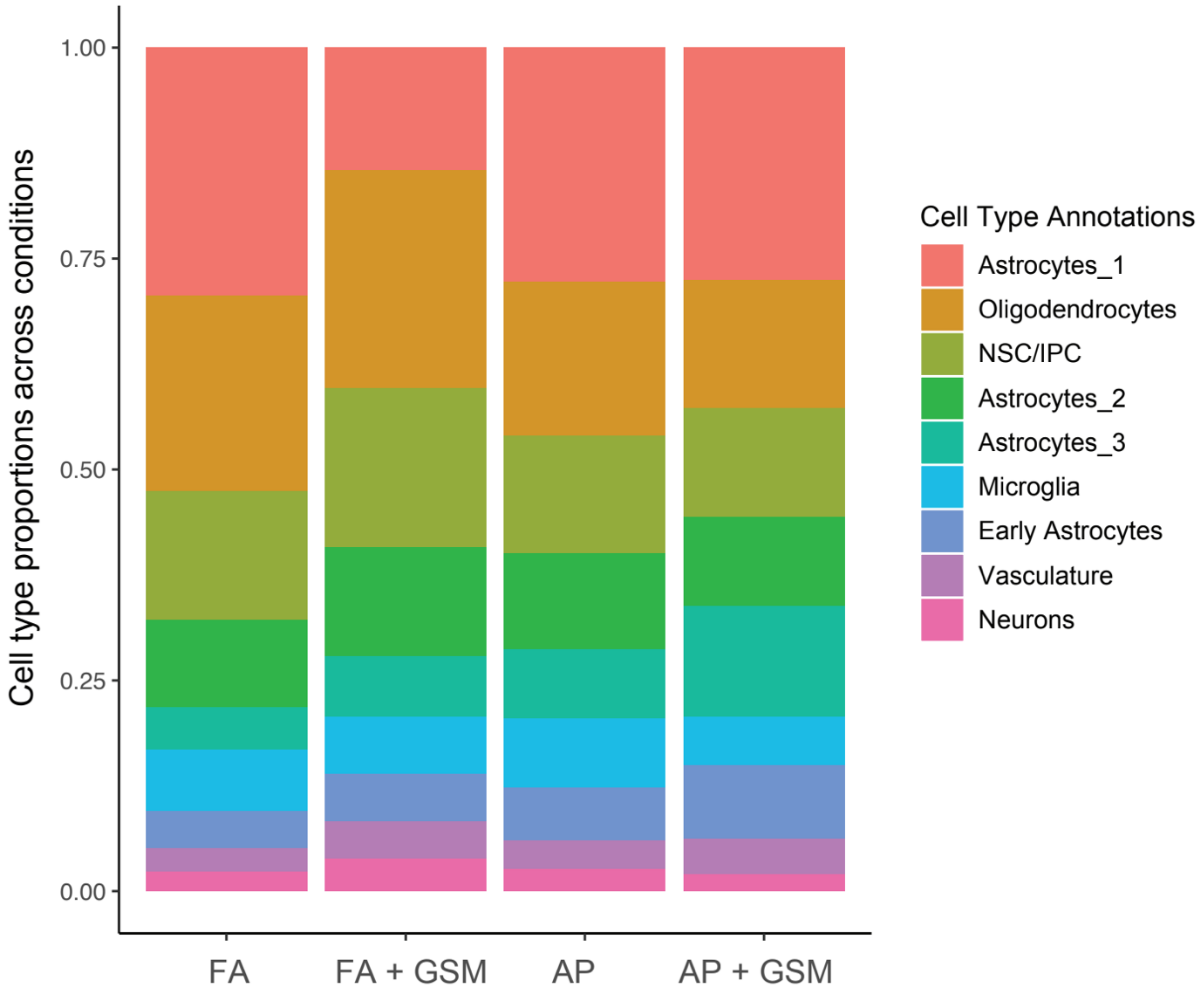


**Supplementary Figure 3: Cell type proportions across conditions**
Stacked bar plot showing the fraction of cells assigned to each annotated cell cluster across experimental conditions.

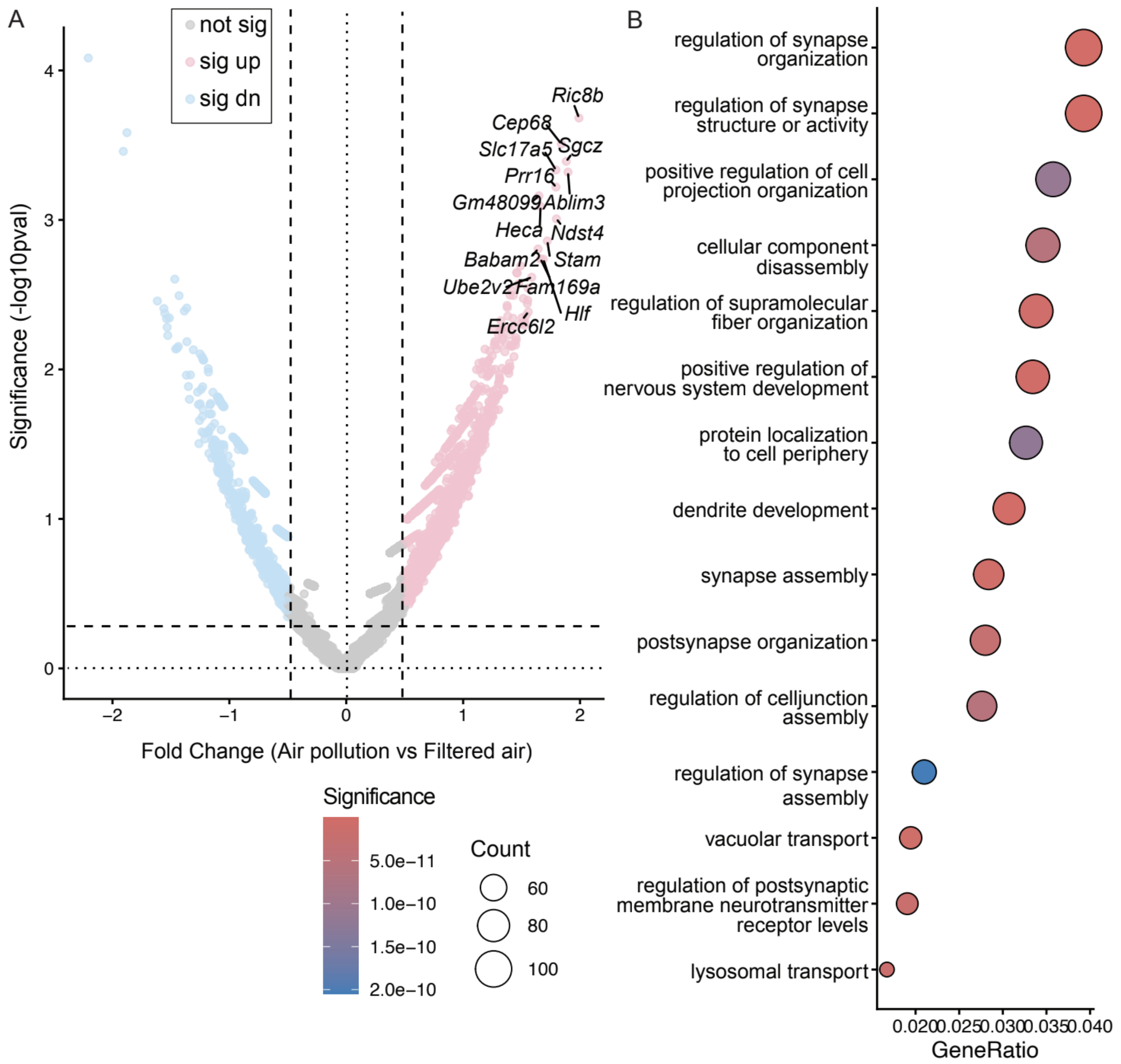


**Supplementary Figure 4. Air pollution induces inflammatory and structural remodeling programs in astrocytes.**

(A) Volcano plot showing differential gene expression in astrocytes comparing air pollution exposure to clean air conditions. The x-axis indicates $\log_2$ fold change, and the y-axis shows statistical significance ($-\log_{10}$ adjusted $p$ value).

(B) Gene ontology enrichment analysis of astrocyte genes significantly upregulated following air pollution exposure. Dot size represents gene count per category, and color denotes adjusted $p$ value. Gene ratio indicates the proportion of differentially expressed genes associated with each term.
